# On some alien species from Emilia-Romagna harbors (northern Adriatic Sea, Italy)

**DOI:** 10.1101/2024.06.13.598796

**Authors:** M. Lezzi, C. Mazziotti

## Abstract

Increased global trade has led to a surge in the introduction of non-indigenous species (NIS) worldwide. Ships, particularly merchant vessels and recreational yachts, act as vectors for these invasions by transporting NIS in ballast water or as hull fouling. This study investigates the introduction of new fouling NIS in the North Adriatic Sea. Our field research at the ports of Ravenna and Cesenatico identified several alien species, including new records for the North Adriatic Sea: *Polyandrocarpa zorritensis, Aplidium accarense* and *Celleporaria brunnea*. Additionally, a significant presence of the cryptogenic species *Molgula manhattensis* was observed in Cesenatico.

## Introduction

The intensification of global trade during recent decades coincides with a substantial rise in the introduction of non-indigenous species (NIS) on a global scale. Vessel traffic acts as a vector for these biological invasions, facilitating the transport of NIS in ballast water or as hull fouling (Katsanevakis *et al*., 2013). Merchant vessels engaged in transoceanic voyages have been identified as the primary facilitators of the introduction of fouling non-indigenous species (Hewitt & Campbell, 2010), alongside recreational yachts that travel regionally and along contiguous coastlines (Ulman *et al*., 2019). However, deliberate introductions also contribute to this phenomenon. Commercially valuable species may be released into novel environments, and farmed specimens can escape from aquaculture facilities. These intentional introductions and transfers within aquaculture represent significant pathways for marine invasions (Katsanevakis *et al*., 2013). The introduction of invasive species does not only occur with the movement of farmed species, but also with farm structures and equipment. Mariculture facilities are located in sheltered coastal spaces, especially estuaries, and provide many artificial hard substrates, thereby offering an optimal habitat for invasive species (Mckindsey *et al*., 2007). So marine aquaculture acts as a source of propagules of invasive fouling species (Lins & Rocha, 2023),

The upper Adriatic Sea is recognized as a hotspot for established non-indigenous species (NIS), for instance are noteworthy the introduction of *Anadara transversa* (Say, 1822), *Anadara kagoshimensis* (Tokunaga, 1906), *Rapana venosa* (Valenciennes, 1846), *Callinectes sapidus* (Rathbun 1896) (Occhipinti-Ambrogi *et al*., 2011). In this study, we report new records of alien species for two port localities on the Romagna coast: port of Ravenna, Italy, and of Cesenatico (North Adriatic Sea).

## Materials and Methods

Sampling activities were conducted at the port of Ravenna, Italy, North Adriatic Sea (44°29’29.7”N 12°17’24.0”E) and of Cesenatico (44°12’21.3”N 12°23’47.5”E) (Fig. 1). Ropes and other submerged objects were randomly observed and collected from the port docks. Additionally, fouling was scraped from the surfaces of floating docks and submerged concrete pylons in some areas. Fouling specimens were collected and passed through a 0.5 mm mesh and fixed with 5% formaldehyde solution. Specimens were preserved in 70% ethanol and identified. Morphological observations and photographic documentation of the specimens were performed using a Nikon DS-Ri2 camera coupled to a Nikon Eclipse e100 microscope for high-resolution imaging and a Nikon SMZ18 stereo microscope for low-magnification analysis. The specimens were deposited in the Laboratory of Benthos Ecology collection of the Oceanographic Unit Daphne of ARPAE, Regional Agency for Environmental Protection and Energy of Emilia Romagna.

**Fig. 1.**
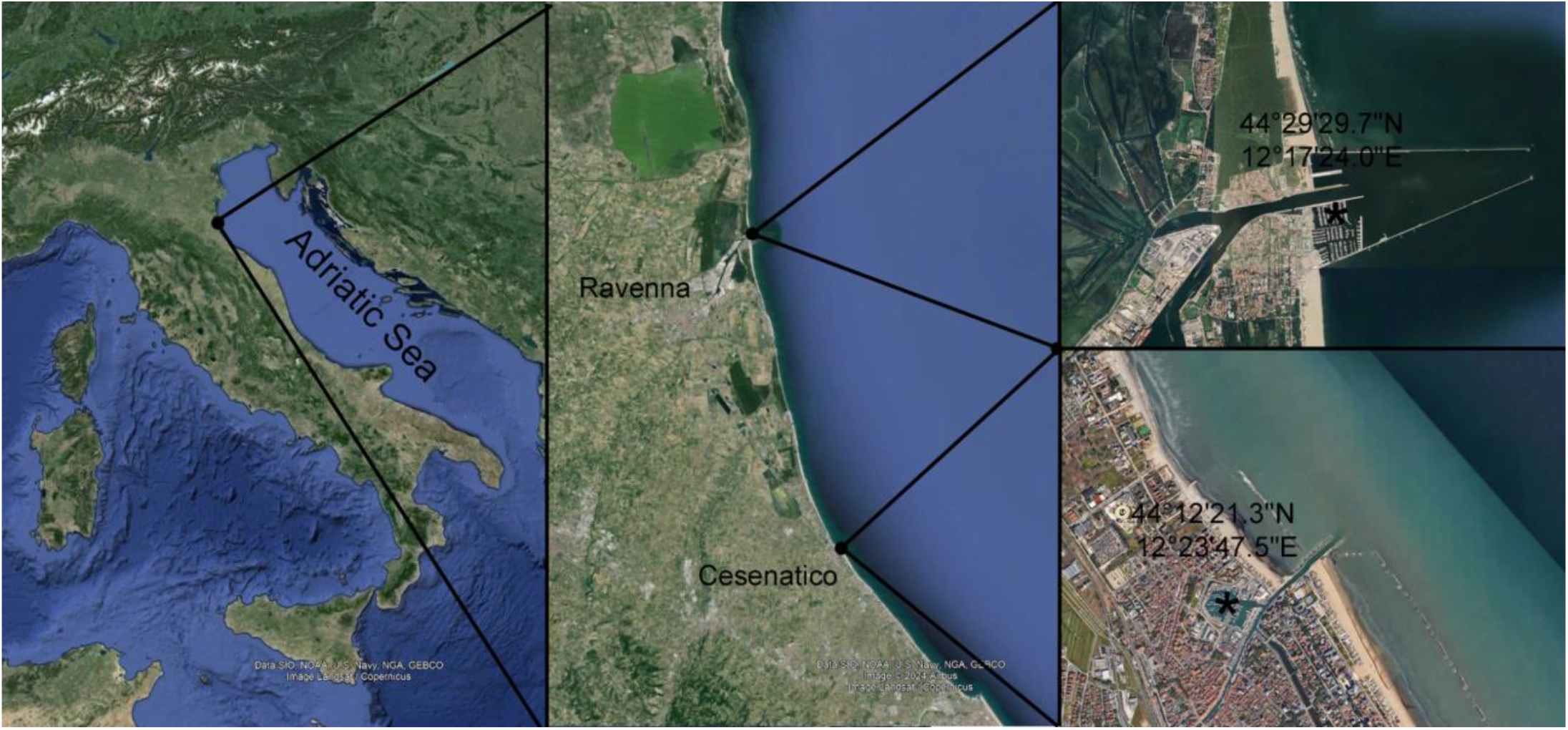
Sampling sites in the North Adriatic Sea: port of Ravenna, Italy, North Adriatic Sea (44°29’29.7”N 12°17’24.0”E) and of Cesenatico (44°12’21.3”N 12°23’47.5”E)

## Results

The material identified included several species known from the North Adriatic sea. Among them, *Caprella scaura* Templeton, 1836, *Paracerceis sculpta* (Holmes, 1904), *Paranthura japonica* Richardson, 1909, *Grandidierella japonica* Stephensen, 1938, *Styela plicata* (Lesueur, 1823), *Amathia verticillata* (delle Chiaje, 1822), *Hydroides elegans* (Haswell, 1883), *Magallana gigas* (Thunberg, 1793), *Pileolaria militaris* Claparède, 1870, *Arcuatula senhousia* (W. H. Benson, 1842), *Amphibalanus amphitrite* (Darwin, 1854) were commonly found in the samples.

Notably, during the summers of 2022 and 2023, the presence of *Polyandrocarpa zorritensis* (Van Name, 1931) and *Aplidium accarense* (Millar, 1953) was recorded in the port of Ravenna, representing new findings for the Adriatic region. Additionally, an unrecorded occurrence of *Celleporaria brunnea* (Hincks, 1884) was documented, representing its first appearance on the Italian side of the North Adriatic coast. Furthermore, a significant presence of the cryptogenic species *Molgula manhattensis* (De Kay, 1843) was reported during the winter of 2019 in the port of Cesenatico.

A brief description and the photographs of these alien species are given below.

### *Polyandrocarpa zorritensis* (Van Name, 1931) (Fig. 2 A, B)

First described from the Peruvian coast, the colonial ascidian *Polyandrocarpa zorritensis* (Stolidobranchia: Styelidae) has emerged as a prominent invasive species in temperate coastal regions in recent decades (Brunetti & Mastrototaro, 2004).

**Fig. 2.**
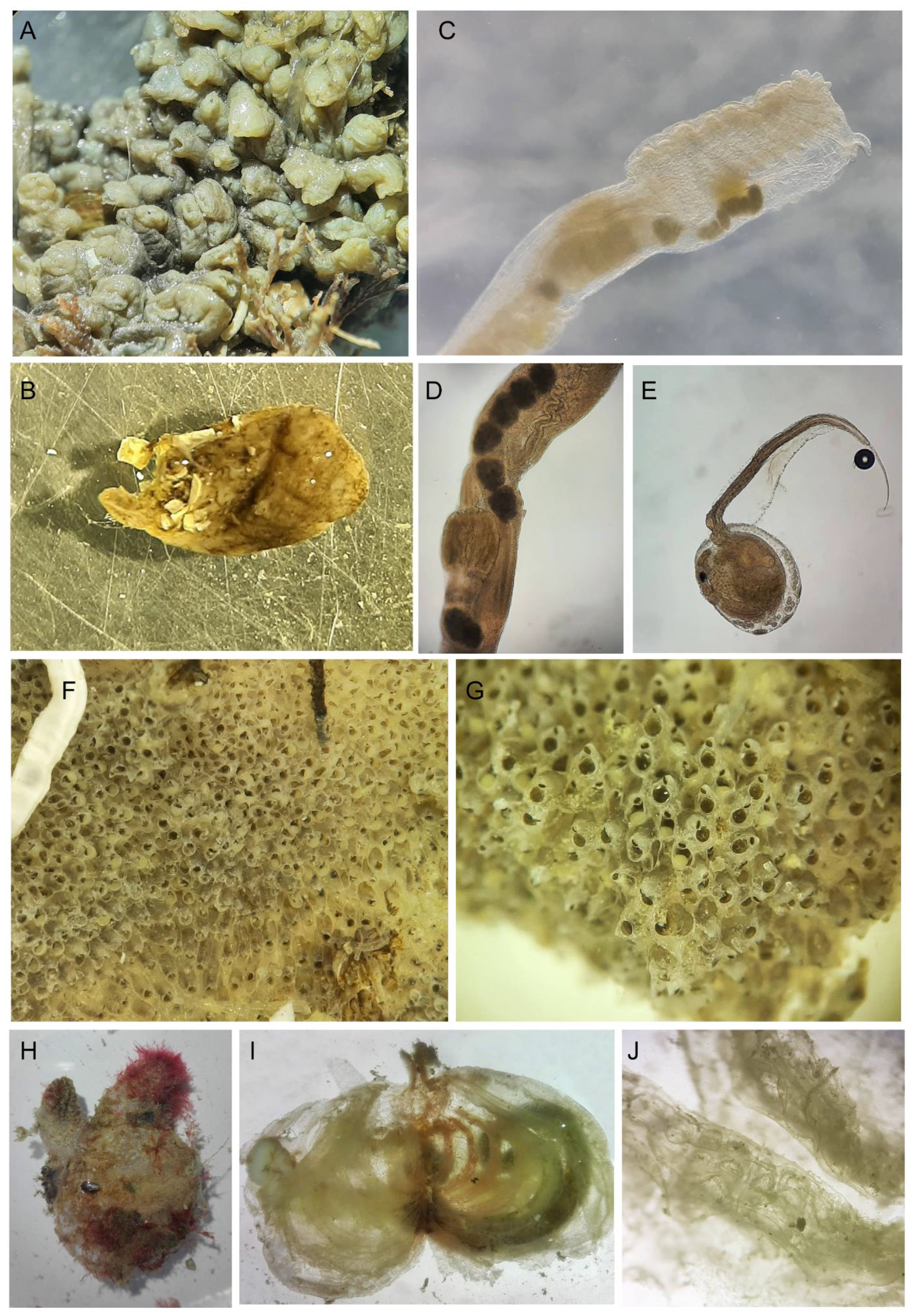
*Polyandrocarpa zorritensis*. A) Colony of *P. zorritensis;* B) Zooid extracted from the colony. *Aplidium accarense:* C) Zooid extracted from the colony showing nine rows of stigmata; D) stomach with the longitudinal fold; E) larvae of *A. accarense. Celleporaria brunnea*: F) Colony; G) zooid evidencing suboral avicularium. *Molgula manhattensis:* H) *M. manhattensis*; I) dissected *M. manhattensis* with branchial sac; J) Magnification of the stigmata showing pustular appearance of the branchiae.

Early records of *P. zorritensis o*utside its native range emerged from the Italian Mediterranean coast, with subsequent reports documenting its establishment in various Mediterranean harbors and gulfs (Brunetti & Mastrototaro, 2004 ; Lezzi *et al*., 2018 ; Tobias-Santos *et al*., 2024). Beyond the Mediterranean, the species has been documented in Pacific and Atlantic regions, including Japan, southern California, Hawaii, the Galapagos Islands, the Panama Canal, the Caribbean, Brazil, the Gulf of Mexico, Florida, and North Carolina (Tobias-Santos *et al*., 2024).

The observed *P. zorritensis* fits with the description of (Brunetti & Mastrototaro, 2004). This species, as evidenced by records from the port of Ravenna, exhibited dominance for the first time during the summer seasons of both August 2002 and July 2023. These temporal observations highlight the recurring nature of its prevalence and suggest a certain degree of consistency in its dominance within the benthic community over the years. The colonies thrive on solid substrates in shallow waters at a depth of 1 meter, amidst *Magallana gigas* specimens.

The colony consists of tightly clustered zooids of varying sizes which are joined by basal stolons. However, these zooids otherwise maintain independence from each other.

The basal part of the colony forms a complex network of stolons, adorned with orange globular bodies (buds) that will eventually develop into new zooids. Fully developed zooids exhibit a sub-cylindrical shape, displaying a yellow-green coloration, and feature an apical oral siphon along with a slightly off-center atrial siphon. Each aperture is characterized by four lobes, each adorned with two distinct dark, almost black, bands.

Well-developed zooids range in height from 10 to 12mm. The body wall has delicate musculature with regularly arranged longitudinal and transverse fibers of the same size. The branchial sac is characterized by four slender, low folds on each side, hosting up to 12 internal longitudinal vessels along the folds and up to two vessels in between.

Between twenty to thirty simple tentacles are present that come in two different lengths. The dorsal lamina is flat with a smooth edge. This species, as noted in various port areas, has the propensity to emerge as the dominant one in the local benthic community during the summer seasons (Lezzi *et al*., 2018). The observed prevalence of this species underscores its ecological significance, suggesting a notable impact on the composition and dynamics of the benthic community in port environments. The heightened dominance during summer may be attributed to favorable environmental conditions, such as temperature, nutrient availability, or other ecological factors that facilitate its thriving in this specific period.

While locally abundant, *P. zorritensis* does not rank among the most widespread non-indigenous ascidians in the Mediterranean (Ulman *et al*., 2019). For example, the species was only found few marinas surveyed by both (Ferrario *et al*., 2017), (Ulman *et al*., 2019) and (López-Legentil *et al*., 2015).

Therefore, despite its long-standing presence in the Mediterranean, the distribution of P. zorritensis has remained relatively confined to the northwestern basins (Tobias-Santos *et al*., 2024). This record from the northern Adriatic region represents the first documented expansion of the species, highlighting its potential as an expanding invasive, particularly within marinas, suggesting hull fouling as a possible dispersal vector.

### *Aplidium accarense* (Millar, 1953) (Fig. 2 C, D, E)

*Aplidium accarense* (Millar, 1953), an aplousobranch ascidian, was initially described along the western coast of Africa, being recognized as native to that region. Recently, *A. accarense* has been observed along the Catalan coasts and in the Tyrrhenian Seas of the Western Mediterranean, as well as in Taranto (López-Legentil *et al*., 2015 ; Montesanto *et al*., 2021). In these locations, it is identified as a non-indigenous species, with previous records limited to harbors and aquaculture farms. The recent recording of *A. accarense* in the harbor of Ravenna, observed during the summer of 2023, provides evidence supporting the idea that the species is actively spreading within the surrounding basin. This observation raises the possibility that *A. accarense* might exhibit an invasive tendency in the local area, posing potential ecological implications for the ecosystem. *Aplidium accarense* was identified according to Montesanto et al. (2021).

*Aplidium accarense* presents as encrusting colonies with a yellowish hue and a slightly cushion-shaped structure, typically measuring around 2 cm in thickness. The tunic of this ascidian is soft and translucent, devoid of embedded sand.

The colonies of *A. accarense* exhibit unrestricted growth, often intermingling with bryozoans and other ascidians such as *Styela plicata*. The individual zooids within the colonies measure up to 10 mm in length, featuring a notably elongated post-abdomen, which can extend up to 6 mm. Each zooid possesses a lobed oral siphon, characterized by six small pointed lobes. Additionally, the atrial siphon is equipped with a short, simple languet emerging from the upper rim, situated at the level of the first row of stigmata.

The thorax is notably longer than the abdomen. The pharynx has 7-10 rows of stigmata in the observed zooid, featuring approximately 10-11 stigmata per half row. Within the abdomen, the alimentary canal houses a plicated stomach exhibiting around 15-17 longitudinal folds, occasionally broken. Following the stomach, the intestine undergoes twisting, forming a subsequent dilation of the gut loop. The digestive system includes the presence of two proximal caeca at the base of the rectum, which terminates in an anus positioned midway up the branchial sac, precisely at the level of the 4th stigmata row.

Observations of larvae reveal distinctive characteristics in the anterior end, including three adhesive organs and several spherical ectodermal ampullae arranged in four groups of 5-8.

### *Molgula manhattensis* (De Kay, 1843) (Fig. 2 H, I, J)

*Molgula manhattensis* (De Kay, 1843), also known as Sea Grape, is a solitary tunicate with rounded and globular shape, which was first described from New York harbour. It has a North-west Atlantic distribution that extends from Massachusetts to southern Louisiana (Van Name, 1945). The European North-east Atlantic distribution of *M. manhattensis* is patchy extending from Norway to Portugal (Monniot, 1969). *Molgula manhattensis* has a recent history of world-wide introductions (Haydar *et al*., 2011).

Hydar et al. 2011, investigating on the native range of *M. manhattensis* by using the molecular approach, concludes that the species is probably native from North-east America, even though its status remains unresolved in European waters and additional molecular studies focused on haplotype distribution remain necessary (i.e. no studied were conducted on Mediterranean populations). Hydar et al. 2011, considered *M. manhattensis* cryptogenic in Europe.

We reported, after the first record made by (Monniot, 1969), the cryptogenic *M. manhattensis* in the north Adriatic. Several specimens were sampled as foulers of *M. giga*s in the harbour of Cesenatico. Grapes of about 3-4 specimens were present on a single *M. gigas* shell and along ropes.

The specimens align with the redescription provided by (Monniot, 1969) for *M. manhattensis*. Externally, the species displays a circular to oval shape, appearing white to translucent, and is covered with short hairs and sand or muddy particles. Internally, the analyzed specimens exhibit distinct features. The dorsal side shows a U-shaped slit with a short dorsal lamina, forming a dorsal tubercle. The branchial sac is characterized by six narrow folds, each containing five internal broad longitudinal vessels. Within these folds, infundibula are divided into four, and distinctive exo-infundibula on the ventral side contributing to a pustular appearance of the branchiae. Papillae on the branchial wall are also evident. The gut loop appears narrow and curved, and the left gonad runs parallel to the kidney. Simultaneously, the right gonad is found within the gut loop.

### *Celleporaria brunnea* (Hincks, 1884) (Fig. 2 F, G)

*Celleporaria brunnea* (Hincks, 1884) is a bryozoan species naturally occurring in fouling communities along the North-east Pacific coasts (Soule, 1961). Recently, *C. brunnea* has been identified beyond its native range, with documented occurrences along the Portuguese and French Atlantic coasts and in South Korea. Additionally, *C. brunnea* has been reported in the Eastern Mediterranean Sea, specifically along the coasts of Turkey, Lebanon, Italy (Lezzi *et al*., 2015 ; Lodola *et al*., 2015)

For the first time, we report the presence of *C. brunnea* in the Italian side of the North Adriatic Sea. This discovery provides evidence of the further spread of *C. brunnea* along the Italian coast, particularly in harbor environments.

Specimens of *C. brunnea* were observed on both *M. gigas* and the ropes. The colonies exhibit an encrusting growth pattern and display a grey to light-brown coloration. The zoecia are irregularly arranged, with a smooth surface more or less punctuated at the margin. The orifice is characterized by a rounded pseudosinus in the proximal margin, delineated by two denticular projections. There are two oral spines, accompanied by a brown basal joint that may be lost and concealed through calcification. Three or, rarely, four spines are present, typically near the peripheral budding margin.

The peristome is raised perpendicularly to the frontal plane, supporting an avicularium on one side, positioned close to the top, and featuring denticulations in the distal rim. Large avicularia are rare and exhibit a spatulate mandible. Both mandibles and opercula are dark brown in color.

## Conclusion

The detailed study occurred in the the Harbour of Ravenna and Cesneatico, in addition to confirming that the presence of non-indigenous species already frequently reported in the fouling communities of the northern Adriatic, contributes to a better knowledge with new records in the examined ecoregion. These findings add an important dimension to our understanding of the ecological dynamics in the region, emphasizing the need for continued monitoring and research to comprehend the impact of alien species on local marine ecosystems.

Furthermore, this discovery underscores the North Adriatic’s vulnerability to invasive species. The intense maritime traffic, a well-known vector for introducing and spreading such species throughout the Mediterranean Sea (Galil, 2006 ; Galil, 2012) , coupled with the physical characteristics of the Adriatic Sea, including its numerous commercial and tourist ports, marinas, and aquaculture sites, creates ideal conditions for colonization by invasive species (Spagnolo *et al*., 2019).

Specifically, enclosed harbors like Ravenna and Cesenatico are susceptible to poor water quality due to several factors such as urban runoff, industrial wastewater discharge, limited tidal flow, and reduced exposure to wind and waves. These factors create a clear environmental gradient, altering salinity and pollutant concentrations, which significantly influence the establishment of non-indigenous species (NIS) (López-Legentil *et al*., 2015 ; Airoldi *et al*., 2015) and this occurrence further emphasizes the vulnerability of such environment to the invasion of alien species.

